# Intragenus (*Homo*) Variation in a Chemokine Receptor Gene (*CCR5*)

**DOI:** 10.1101/195081

**Authors:** Kara C. Hoover

## Abstract

Humans have a comparatively higher rate of more polymorphisms in regulatory regions of the primate *CCR5* gene, an immune system gene with both general and specific functions. This has been interpreted as allowing flexibility and diversity of gene expression in response to varying disease loads. A broad expression repertoire is useful to humans—the only globally distributed primate—due to our unique adaptive pattern that increased pathogen exposure and disease loads (e.g., sedentism, subsistence practices). The main objective of the study was to determine if the previously observed human pattern of increased variation extended to other members of our genus, *Homo*. The data for this study are mined from the published genomes of extinct hominins (four Neandertals and two Denisovans), an ancient human (Ust’-Ishim), and modern humans (1000 Genomes). An average of 15 polymorphisms per individual were found in human populations (with a total of 262 polymorphisms). There were 94 polymorphisms identified across extinct *Homo* (an average of 13 per individual) with 41 previously observed in modern humans and 53 novel polymorphisms (32 in Denisova and 21 in Neandertal). Neither the frequency nor distribution of polymorphisms across gene regions exhibit significant differences within the genus *Homo*. Thus, humans are not unique with regards to the increased frequency of regulatory polymorphisms and the evolution of variation patterns across *CCR5* gene appears to have originated within the genus. A broader evolutionary perspective on regulatory flexibility may be that it provided an advantage during the transition to confrontational foraging (and later hunting) that altered human-environment interaction as well as during migration to Eurasia and encounters with novel pathogens.

## Introduction

Chemokine receptors facilitate communication between cells and the environment [1, 2] and mediate the activity of chemokines, proteins secreted by the immune system genes to chemically recruit immune cells to infection sites via chemotaxis [2, 3]. The cell surface chemokine receptor *CCR5* (a G protein-coupled receptor) is best known for its adaptive immune system role in binding the M-tropic human immunodeficiency virus (HIV) and creating a gateway to host cell infection [3–12]. In several mammals, *CCR5* genes present high levels of gene conversion with the chromosomally adjacent *CCR2* [13–18]. Primate *CCR5* gene structure, open reading frame (ORF), and amino acid identity are evolutionary highly conserved [2, 19–21] and interspecific gene sequences are functionally similar [20]. There is, however, common and significant variation across species outside conserved regions. Most of these polymorphisms are not deleterious to health and tolerated due to the redundancy of the chemokine family in ligand binding [2, 22]. New World Monkeys have a high number of functional polymorphisms due to lentivirus resistance [23]. Further, humans have been found to have a significantly high number of *cis*-regulatory region polymorphisms in comparison to 36 non-human primate species of apes, Old World Monkeys, and New World Monkeys [20]. Humans also have a specific a 32bp deletion in Exon 3, *CCR5Δ32* [24, 25], that results in a non-functional protein [24–29]) associated with HIV-resistance and West Nile Virus susceptibility in northern European populations [30–42].

Located on Chromosome 3 (3p21), human *CCR5* is 6,065 bases long with an ORF of 1,056 bases that codes for a protein with 352 residues. Two common transcripts (B with three exons and the more stable A with four) likely resulted from non-coding upstream polymorphisms in two separate gene promotors (the functionally weaker *cis*-acting promoter (P_U_) upstream of Exon 1 and the downstream promoter (P_D_) upstream of Exon 3 [19, 20]. These transcripts cause alternate splicing (differential inclusion or exclusion of exons) in messenger RNA that affects regulation of cell surface receptor expression levels [20–22].

The plasticity in regulation of gene expression via alternate transcripts and increased polymorphisms [20–22] makes human *CCR5* particularly interesting from a broader evolutionary perspective. *Homo* has one of the broadest adaptive ranges of any species [43] and human CCR5’s ability to rapidly respond to new pathogens [19, 20] may have served an adaptive function during evolutionary migration and shifts in human-environment relationships with changes to subsistence. Our genomes carry vast evidence of past disease responses [44–46] that are shared across the genus and reflect a unique disease pattern for *Homo*. For example, there is strong evidence for increased disease risk via genetic load in extinct *Homo* and past human populations [47] and archaeological evidence for past disease treatment (ingestion of anti-biotic and anti-fungal non-food plants) in Neandertals [48–52]. *CCR5* has been well studied due to its role in HIV infection (with a focus on natural selection acting on the 32bp deletion) but no work has explored variation within the genus *Homo* more broadly.

The plethora of research on the evolution *CCR5* was conducted prior to the generation of deep coverage, high quality paleogenomes for extinct hominins, such as Neandertal species and the newer Denisova species. While paleogenomic sample sizes are not robust to make statements on selection or add to a discussion of other evolutionary forces acting on variation, they provide an evolutionary dimension to understanding the patterns of variation characterizing our genus and insights into possible adaptations to new environments, subsistence regimes, and pathogens [53]. Plus, the sample of ancient genome is increasing every year. Just a few insights gained from a single paleogenomes include ground-breaking studies on evolution of skin color in humans [54]and Neandertals [55] and the introgression of functionally adaptive polymorphisms into the human immune system genes from Altai Neandertal [56]. Understanding the differences between derived and specific variation also enables potential differentiation of challenges we overcame as a genus such as obligate bipedalism [57] or high-altitude adaptation [58] versus challenges we overcame as a species such as the biocultural evolution of sickle-cell trait and malaria infection [59]. Thus, the overall aim of this research is to place humans within the context *Homo* and examine if the pattern of humans having a significantly higher number of *cis*-regulatory region polymorphisms (compared to 36 non-human primate species of apes, Old World Monkeys, and New World Monkeys) [20] is specific or one that is shared by our genus.

Variation in *CCR5* was examined in humans and extinct hominins to address the questions: are there shared patterns of variation across the genus for polymorphism frequency and is the distribution of polymorphisms across the gene suggestive of a common evolutionary trajectory? Based on previous studies on human-nonhuman primate gene structure and variation and the finding that human polymorphisms allow flexible *CCR5* gene expression [19, 20], the expectation is that there is a shared pattern of variation that aided adaptation for members of the geographically and ecological dispersed genus *Homo*. Both expectations were met.

## Materials

Modern human data is from the 1000 Genomes Project [60], which contains data for 2,504 individuals from 26 populations (Table 1). While coverage is low per individual, the data are robust enough to identify the majority of polymorphisms at a frequency of at least 1% in the populations studied, which is suitable for the current study. Extinct *Homo* data are from two Denisova samples (Denisova 3 and 2) and four samples from the Neandertal Genome Project (Vindija, Altai, El Sidron, Mezmaiskaya). These species are Pleistocene Eurasian hominins with Denisova representing an eastern Eurasian Pleistocene population and Neandertal a western one (with some overlap with Denisova in Siberia). All genomes have high coverage (excepting Mezmaiskaya and Sidron); contamination with modern human DNA is estimated to be less than 1% for the extinct hominins [50, 61–64].

**Table 1:** Samples plus polymorphism totals

| Region | Population Name | n | Var | AvVar/Pop |
| --- | --- | --- | --- | --- |
| Africa | African Caribbeans in Barbados | 96 | 1464 | 15 |
|  | Americans of African Ancestry in SW USA | 61 | 1049 | 17 |
|  | Esan in Nigeria | 99 | 1499 | 15 |
|  | Gambian in Western Divisions in the Gambia | 113 | 1744 | 15 |
|  | Luhya in Webuye, Kenya | 99 | 1469 | 15 |
|  | Mende in Sierra Leone | 85 | 1420 | 17 |
|  | Yoruba in Ibadan, Nigeria | 108 | 1786 | 17 |
| America | Colombians from Medellin, Colombia | 94 | 1688 | 18 |
|  | Mexican Ancestry from Los Angeles USA | 64 | 1111 | 17 |
|  | Peruvians from Lima, Peru | 85 | 1184 | 14 |
|  | Puerto Ricans from Puerto Rico | 104 | 1699 | 16 |
| East Asia | Chinese Dai in Xishuangbanna, China | 93 | 2222 | 24 |
|  | Han Chinese in Beijing, China | 103 | 2278 | 22 |
|  | Southern Han Chinese | 105 | 2328 | 22 |
|  | Japanese in Tokyo, Japan | 104 | 1929 | 19 |
|  | Kinh in Ho Chi Minh City, Vietnam | 99 | 2411 | 24 |
| Europe | Utah Residents (CEPH) with Northern and Western European Ancestry | 99 | 1664 | 17 |
|  | Finnish in Finland | 99 | 1484 | 15 |
|  | British in England and Scotland | 91 | 1417 | 16 |
|  | Iberian Population in Spain | 107 | 1755 | 16 |
|  | Toscans in Italia | 107 | 2132 | 20 |
| South Asia | Bengali from Bangladesh | 86 | 1729 | 20 |
|  | Gujarati Indian from Houston, Texas | 103 | 1818 | 18 |
|  | Indian Telugu from the UK | 102 | 1907 | 19 |
|  | Punjabi from Lahore, Pakistan | 96 | 1831 | 19 |
|  | Sri Lankan Tamil from the UK | 102 | 1952 | 19 |
| *Homo* | Altai | 1 | 7 | 7 |
|  | Denisova 3 | 1 | 10 | 10 |
|  | Denisova 2 | 1 | 39 | 39 |
|  | El Sidron | 1 | 0 | 0 |
|  | Mezmaiskaya | 1 | 28 | 28 |
|  | Ust'-Ishim | 1 | 15 | 15 |
|  | Vindija | 1 | 8 | 8 |

## Methods

The human reference sequences for two common transcript variants for the *CCR5* gene (NM_000579 and NM_001100168) were downloaded from the National Center for Biotechnology Information (NCBI). The modern human variation data (*CCR5* and *cis*-acting elements) were downloaded from 1000 Genomes via ftp as variant call format (VCF) files (ftp://ftp.1000genomes.ebi.ac.uk/vol1/ftp/). All data were downloaded to and analyzed using the University of Alaska Research Computing Systems. All files were aligned to the human genome GRCh37/hg19. VCF files for six extinct hominin species and one extingt human (Ust’-Ishim) were downloaded from the Max Planck Institute Leipzig. Ancient DNA often contains C-to-T deaminations at the end of reads [for a review see 65]. The lack of variation identified from paleogenomic sequence reads is unlikely to be a result of typical problems associated with ancient DNA sequence reads since chemical processes like deamination would increase SNPs (whether false or not). More significantly, the paleogenomes were generated using protocols [typically as reported in 66] that largely eliminates this error. Despite high levels of variation at this locus and evidence for balancing selection in humans at this locus, strong levels of introgression from inter-breeding with Neandertals in Eurasia have not been reported at this locus, as they have for other immune system loci in similar scenarios [27, 67–72]; introgression data from the Reich lab (https://reich.hms.harvard.edu/datasets) [73] confirm this is the case. Moreover, the African genetic variation is similar to European genetic variation which suggests that diversity was already present in modern humans prior to any admixture with archaic species in Europe.

Distribution of polymorphisms was guided by the structure provided by Mummidi et al. [19] and included promoter regions (P_U_ and P_D_), ORF, and *CCR5*. The target area for P_U_ was the most inclusive range (-1976 to +33) which avoided overlap with P_D_ and because little difference was noted between putative Pu regions studied by Mummidi et al [19]. The target area for Pd was the most productive range (+119 to +828). Significant difference in distribution of polymorphisms across gene structure for all samples was tested using Monte Carlo methods for the exact test.

## Results and Discussion

Previous research has examined gene structure [19], gene variation within primate species [20], and selection acting on the gene primarily in response to viral load [21, 22, 33, 40, 41, 74] [6, 10, 21, 33, 36, 37, 40, 41, 74, 75]. The goal of this research was to establish if the pattern of human variation and distribution of polymorphisms in *CCR5* [20] is specific (i.e., unique in the human species) or genus-wide (i.e., a pattern shared by *Homo*).

Are there shared patterns of variation across the genus for polymorphism frequency? In the modern human sample, 262 known SNPs were observed (Table 2, Supplementary Table 2 contains all 1000 Genomes variants). SNP frequency per individual (total SNPs in a population/total number of individuals) within the 26 populations ranged from 14 to 24, with East Asians exhibiting the highest variation and Africa and the Americas the least (Table 1). There were 94 polymorphisms identified across all extinct *Homo* samples (Altai, Denisova 3 and 2, Mezmaiskaya, Ust’-Ishim, Vindija, and El Sid), an average of 13 per individual included in analysis (Table 1 and 2; Supplementary Table 1). No polymorphisms were found in the El Sidron specimen and, as a result, it is not included in the tables. Some polymorphisms in extinct *Homo* (n=41) have been previously observed in modern humans (Table 2). There were 53 novel polymorphisms identified, 32 in Denisova (1 in Denisova 3, 31 in Denisova 2) and 21 in Mezmaiskaya. Table 2 summarizes extinct *Homo* polymorphisms.

**Table 2:**
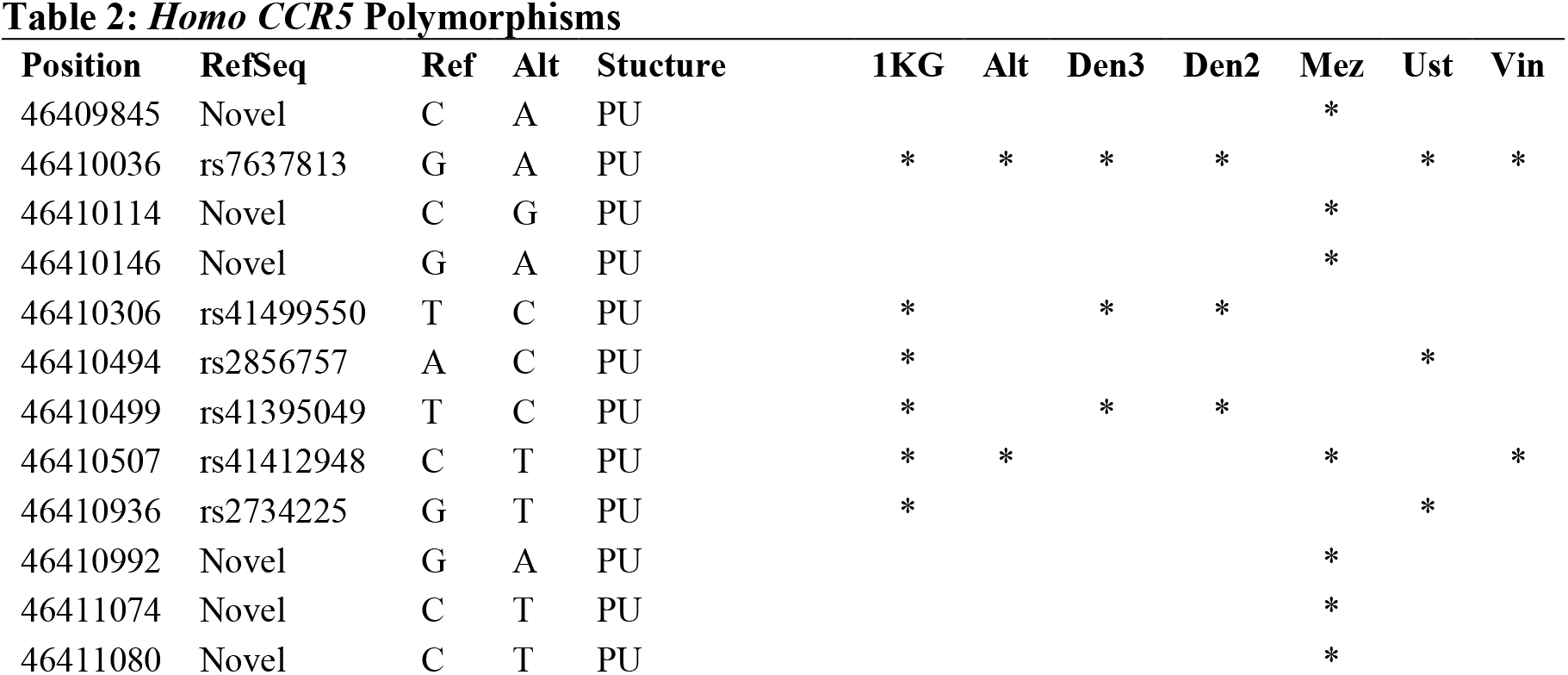
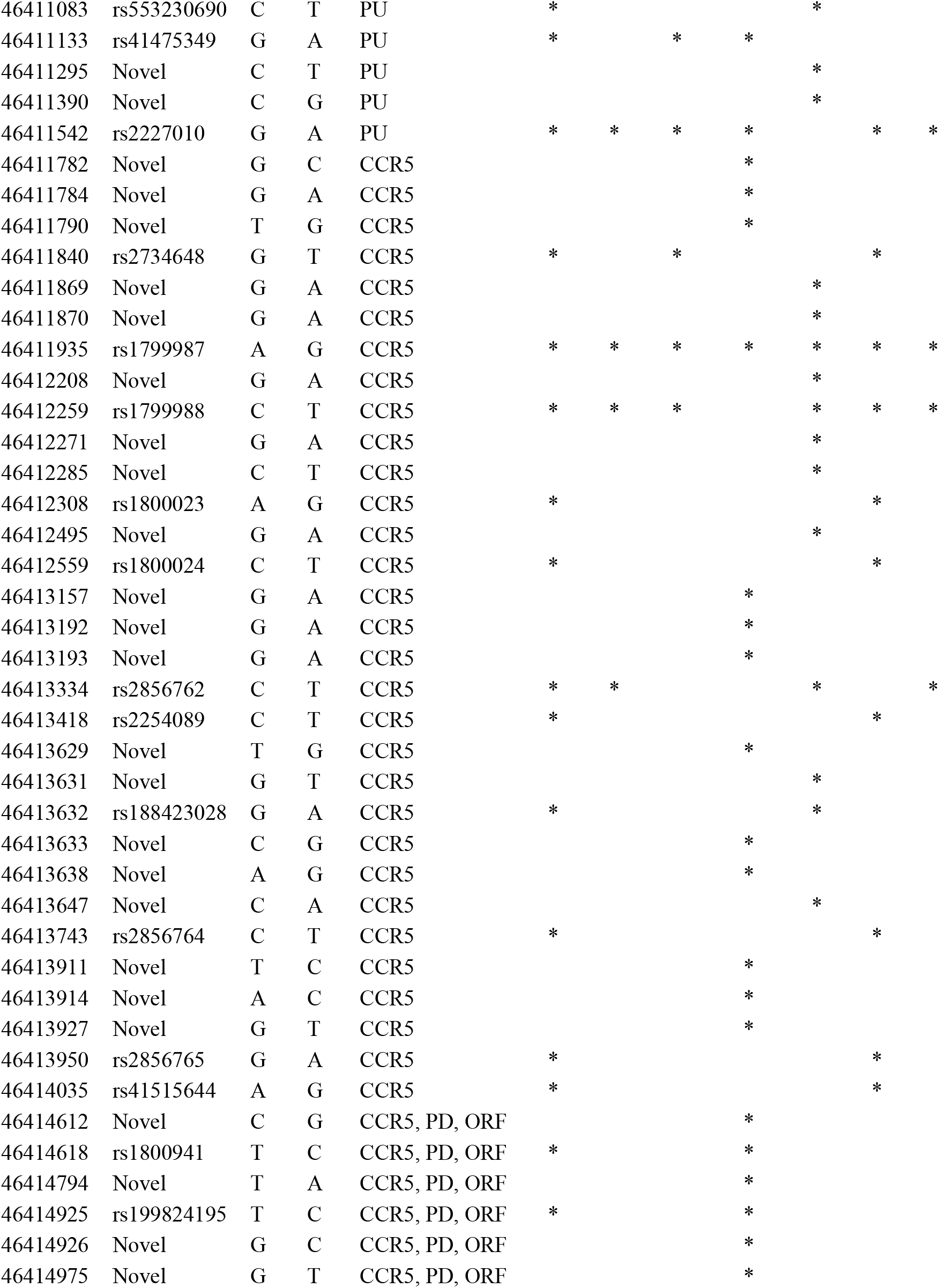
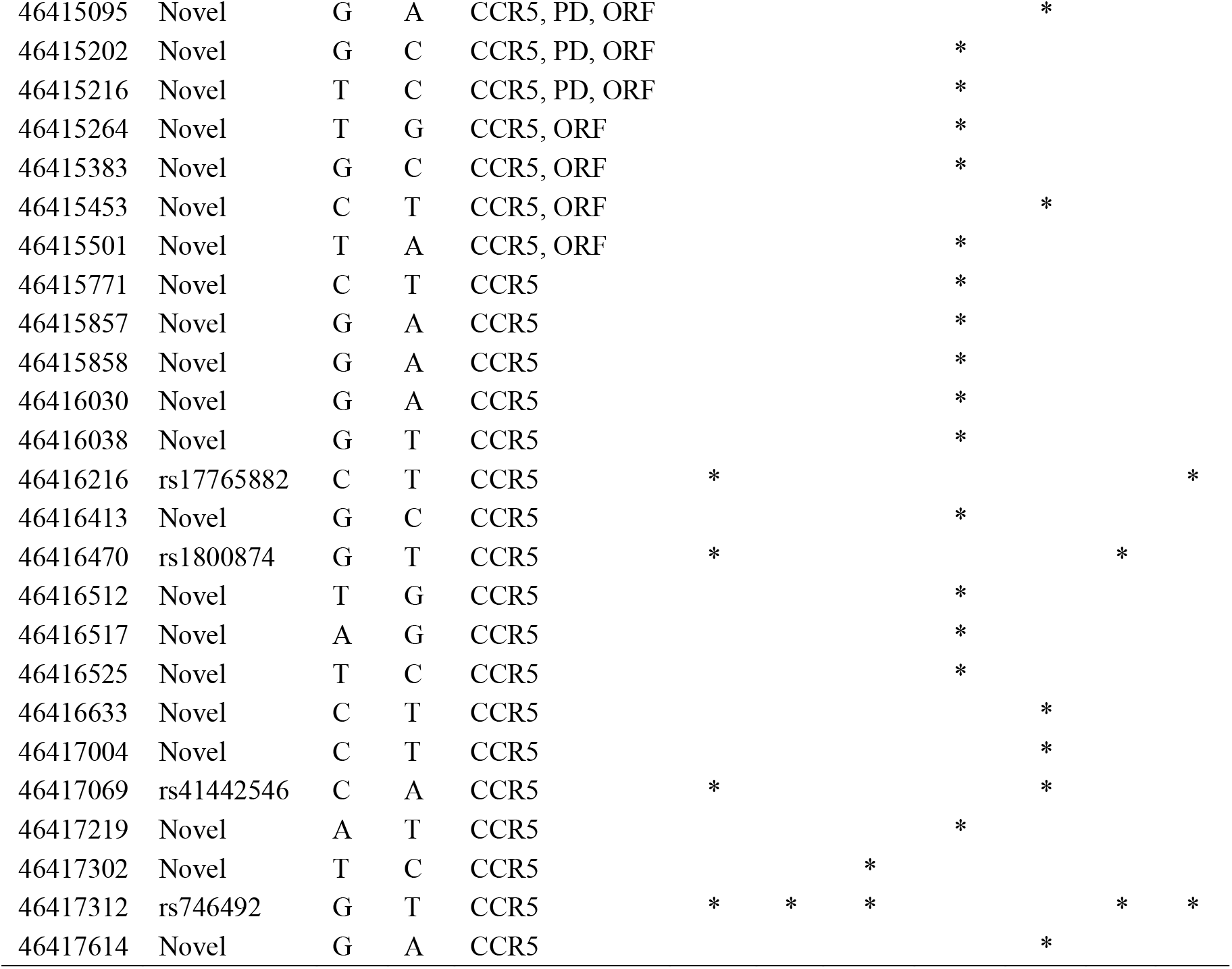
*Homo CCR5* Polymorphisms

Is the distribution of polymorphisms across the gene suggestive of a common evolutionary trajectory? The frequency of polymorphisms across gene structure are used rather than counts because the human sample is much larger and captures an exponentially greater number of polymorphisms as a result (see Supplementary Table 3 for raw count sumamry). Both humans and extinct members of our genus exhibit more polymorphisms in gene regulatory regions (Table 3) suggesting a shared pattern of variation across *Homo*. When polymorphisms occur in both P_U_ and P_D_, there is a greater frequency in the functionally stronger regulatory area, P_D_, but in four ancient samples (Altai, Denisova 3, Vindija, and Ust-Ishim), they only occur in P_U_ (see Supplementary Table 3); only Denisova 2 and Mezmaiskaya had no polymorphisms in the ORF. The comparatively lower frequency across all samples reflects the conservation trend noted in primates [19, 20]. A structural analysis of the distribution of polymorphisms via an Exact Test indicated no significant statistical differences among all samples (results not shown). Given the expected frequency of polymorphism (based on the perception of CCR5 covered by an area of interest—see Table 3 footnotes), there is a significant pattern in the samples. First, modern humans and Denisova 2 have a greater than expected number of polymorphisms in the ORF (even if these are exceeded by polymorphisms in regulatory regions). All samples (except Denisova 2) have a greater than expected number of polymorphisms in the promoter regions.

**Table 3:** Variation in gene structure, polymorphisms as percentage

|  | **Expected** | **Actual** |  |  |  |  |  |  |
| --- | --- | --- | --- | --- | --- | --- | --- | --- |
|  | **RefSeq** | **1kg** | **alt** | **den3** | **den2** | **mez** | **ust** | **vin** |
| ORF | 17.46% | 22.33% | 0.00% | 0.00% | 32.35% | 11.11% | 0.00% | 0.00% |
| P<sub>U</sub> | 33.80% | 32.94% | 75.00% | 100.00% | 28.89% | 55.00% | 36.36% | 60.00% |

<sup>1</sup> Percent CCR5 covered by ORF
<sup>2</sup> Percent Promoter+CCR5 covered by promoters

Prior research found that humans have a potentially unique plasticity in gene expression due to the effect of alternate splicing [57]. The distribution of polymorphisms across gene regions in *Homo* suggests plasticity in gene regulation and expression in response to viral loads, as noted in previous studies [19, 20]. The pattern of immune gene introgression, particularly regulatory haplotypes in the antiviral OAS gene cluster [67], has suggested that selective forces in our close relatives operated on expression, not protein variation—same as seen in non-human primate *CCR5* variation [19, 20]—and those adaptations were also useful to humans. Thus, an increase in polymorphisms that allowed plasticity in regulation and expression in *CCR5* makes sense even if it is not due to introgression. Without functional testing, the exact nature of the polymorphisms is not known other than by inference and comparative analysis, as done here. And, without more paleo-genomes to compare, we cannot know if the variation in these genomes represents true species variation but the data presented here indicate that the pattern is not human specific, rather one shared by recent members of *Homo*.

The expectation that extinct hominins and modern humans would share this pattern of increased variation in the regulatory areas of the gene is met in the current study. Our genus has several unique behavioral and genetic adaptations compared to nonhuman primates and these adaptations might hold some avenues for further research. For instance, a genus-wide shift in subsistence activities occurred during the Plio-Pleistocene (roughly 2 million years ago) from opportunistic non-confrontational scavenging to confrontational scavenging and, later, top predatory behaviors; this alteration to hominin-environment interaction brought hominins into greater and regular contact with animal carcasses [76–81]. Neandertals in Europe have also been shown to be active hunters and foragers [82] who experienced increased pathogen exposure and disease load as a result [48, 49, 51, 83]. Humans and European Neandertals would have shared similar ecological adaptive pressures in Europe—broad and varied— whereas Altai Neandertal (related to European Neandertals) and Denisova would have shared similar ecological adaptive pressures in Siberia with Ust’-Ishim—less varied. Key pathogens year-round in tropical to temperate zones are more likely to be viral (vector-borne) or bacterial (zoonotic) with transmission via interaction with the environment [84]; high latitude pathogens year-round are more likely parasitic due to the reliance on marine mammals [85] and the short season for viral vector reproductive cycles to transmit infection from insects to hominins [35]. Evidence for gene introgression from extinct hominin species to modern humans is clustered (among other domains) in immune system genes [27, 69, 73, 86]; in particular, the *OAS* anti-viral gene cluster on Chromosome 12 shows signatures of positive selection [69, 73], which suggests that adaptation to Eurasian pathogens may have been partly facilitated by prior adaptative mutations to local viral loads. At a minimum, the environmental challenge faced by non-human members of *Homo* facilitated human adaptation to a new environment—a shared challenge with a similar solution.

While previous studies have examined variation in *CCR5*, particularly *CCR5Δ32* which has a more recent origin [36–42], as a product of more recent human-disease interaction, the widespread pattern of increased variation in the gene across the genus *Homo* identified in this study suggests a potential evolutionary adaptation. A key event distinguishing members of the genus *Homo* from the last common ancestor with Australopithecus was the shift to confrontational scavenging and, later, hunting; this alteration to human-environment interaction added a new point of disease contact as evidenced by modern data showing hunting bushmeat (which ancient hominins did too [80]) alters disease exposure via introduction of retroviruses and other pathogens [87–91]. Given *CCR5*’s role in both innate and adaptive immune system functioning, its plasticity may have provided an advantage to members of *Homo* across these varied disease ecologies and its potentially greater than normal interaction with the environment in foraging and hunting activities. As more ancient genomes become sequenced, we can have more robust data with which to work and invest resources into functional testing and experimentation of what function these polymorphisms might have had.

## Acknowledgments

None.

## Supplementary Information

S1 Table: Altai Neandertal Introgression Data

S2 Table: Variants in 1000 Genomes

S3 Table: Raw counts of variants per sample

